# Generation of human-pig chimeric renal organoids using iPSC technology

**DOI:** 10.1101/2024.03.19.584036

**Authors:** Koki Fujimori, Shuichiro Yamanaka, Kentaro Shimada, Kenji Matsui, Shiho Kawagoe, Takao Kuroda, Atsushi Ikeda, Makoto Inoue, Eiji Kobayashi, Takashi Yokoo

## Abstract

The potential of using porcine organs and human induced pluripotent stem cell (iPSC)-derived organoids as alternative organs for human transplantation has garnered growing attention. However, both approaches still face technological challenges. Interspecies chimeric organ production using human iPSCs is expected to be another promising approach that addresses the challenges associated with organ production. Our research group successfully generated human-mouse chimeric renal organoids by utilizing human iPSC-derived nephron progenitor cells (NPCs) and fetal mouse kidneys. However, the current technology has limited engraftment and development capabilities for human NPCs, and there have been no reports of generating interspecies chimeric renal organoids in larger animals, limited only to rodents. Therefore, in this study, we embarked on the production of human-pig chimeric renal organoids using the pig kidney, which is considered the most promising source of organs for interspecies transplantation to humans. To construct a human-pig chimeric renal organoid culture system, we first modified the existing human-mouse chimeric renal organoid culture system and developed a method that enables the survival and continued renal development of both species. This method was found to be applicable to porcine fetal kidney cells, and ultimately, we successfully produced human-pig chimeric renal organoids. Furthermore, this culture method can also be applied to the generation of human interspecies chimeric kidneys for future clinical applications.

The findings of this study serve as a foundational technology that will greatly accelerate future research in humanized pig kidney production for clinical purposes, and are also expected to be used as an evaluation technique to ensure the quality of human NPCs for xenotransplantation.

## Introduction

In recent years, porcine organs have garnered attention as alternative grafts for human organ transplantation, with several pioneering clinical studies utilizing porcine kidneys having been conducted[1–3]. However, it has become evident that immunological challenges persist in achieving long-term engraftment and functionality[4]. On the other hand, studies on organogenesis utilizing human induced pluripotent stem cells (iPSCs) have also been actively pursued in recent years[5–7]. Nonetheless, the complete establishment of requisite cell types and environments for organogenesis is not yet achieved, necessitating further research to create functional organs[8]. The other method of chimera organogenesis using blastocyst complementation, in which human pluripotent stem cells are injected into embryos genetically engineered not to develop specific organs, has also garnered attention[9–14]. However, there remain ethical issues about the resulting animals, as it is difficult to selectively engraft human cells exclusively into the targeted organs within interspecies organisms[14].

Interspecies chimeric organogenesis by injecting human iPSC-derived cells into developing organs of interspecies animals is envisioned as a promising approach to address these challenges in organ production[15,16]. Indeed, in our previous studies, we achieved the creation of heterologous, including human, chimeric kidneys by utilizing the prenatal kidney development environment of a different species[16,17]. Additionally, our xenotransplantation research demonstrated that heterologous transplantation of porcine fetal kidneys to adult monkeys resulted in lower immune rejection compared to adult organ transplantation[18]. Furthermore, our study using rodents demonstrated that when creating an interspecies chimeric kidney by injecting recipient’s renal cells into the kidney of a different species, the resulting kidney transplanted into another recipient individual has lower immunogenicity compared to conventional interspecies transplantation[19].

Moreover, we are currently engaged in a study aimed at establishing the formation of “chimeric renal organoids” by co-culturing dissociated fetal kidneys and human NPCs at the single-cell level, as a means to enhance the intercellular communication between human and heterologous animal cells, separate from the method for clinical application of injecting human NPCs into heterozygous fetal kidneys. Through this organoid research, we achieved the generation of human-mouse chimeric renal organoids exhibiting accelerated maturation[20].

However, in current techniques, the engraftment and development of human NPCs in both chimeric kidneys and chimeric renal organoids are limited. Furthermore, it has been reported that there are differences in organogenesis rates among interspecies animals[21], and studies using pluripotent stem cells from different species suggest the existence of a barrier preventing interspecies chimerism[22]. These suggest that there are remaining challenges in effectively achieving interspecies chimera formation. Additionally, it remained unclear whether the generation of such chimeric kidneys or chimeric renal organoids could be achieved using human-derived renal progenitor cells in large animals such as pigs, as previous studies were limited to rodent models.

Pigs are considered the most ideal organ xenograft donor, and have several advantages for chimeric kidney production due to their similar organ size, physiological functions resembling humans, rapid growth, and ease of organ procurement[23]. Thus, the generation of chimeric renal organoids using human NPCs and porcine kidney cells represents a crucial step towards the future development of human-pig chimeric kidney organogenesis for clinical application. Nevertheless, no methods for organoid creation using porcine kidney cells or chimeric kidney organoid creation, including the culture of porcine fetal kidney cells, have been reported to date. Furthermore, recent studies employing blastocyst complementation have revealed that human iPSCs and their derivatives are not highly committed to pig blastocysts[14], highlighting the need for establishing cultivation techniques for efficient human-pig chimera formation.

In this study, to construct a human-pig chimeric renal organoid culture system, we initially modified existing human-mouse mixed organoid culture systems to prolong survival and facilitate renal development in their component species. We then validated the applicability of this method to porcine fetal kidney cells, ultimately achieving the creation of human-pig chimeric organoids. Importantly, the identified culture conditions were suitable not only for human-xenogeneic chimeric renal organoids but also for human-interspecies chimeric kidneys for future clinical purposes.

## Results

### Optimization of culture conditions for chimeric renal organoid composition

The production of fetal mouse kidney organoids, in which fetal kidneys are enzymatically processed, aggregated from a single-cell state, and subsequently reconstructed to recapitulate renal structures, has been documented in previous studies[24,25]. Leveraging the self-organization of fetal kidney progenitor cells, we demonstrated the incorporation of human iPSC-derived NPCs during fetal kidney organoid formation, resulting in the formation of interspecies human-mouse chimeric nephrons[20]. However, while we initially considered the survival and organoid formation of cell types constituting the kidney organoids to be crucial, it became evident in our approach that the survival and organoid formation of human NPCs were not sufficient (SupFig. 1b) even when utilizing human NPCs with confirmed differentiation potential (SupFig. 1a). Consequently, it was confirmed that the proportion of human cells within chimeric kidney organoids and the chimera formation rate diminished in a time-dependent manner (SupFig. 1c, d).

Therefore, we further explored culture conditions that would enable the survival, differentiation, and maturation of both chimeric kidney organoids and their constituent cells. We initiated an investigation into the culture conditions using human-mouse chimeric kidney organoids as a starting point (Fig. 1a). For the reaggregation medium and the maturation medium during chimeric kidney organoid formation, we selected previously employed media for differentiation and maturation of NPCs, specifically “NPC_Re-agg” medium for the reaggregation medium, and “NPC_Mat” medium and “KR5_Mat” medium[15,16] for the maturation medium[26–29] (Fig. 1a). Initially, using DMEM/F12 with 10% FBS for reaggregation and either NPC_Mat or KR5_Mat for maturation, organoid formation was observed in all species/condition; however, the reaggregated products from human NPC did not exhibit a distinct organoid morphology (Fig. 1b).

**Fig. 1.**
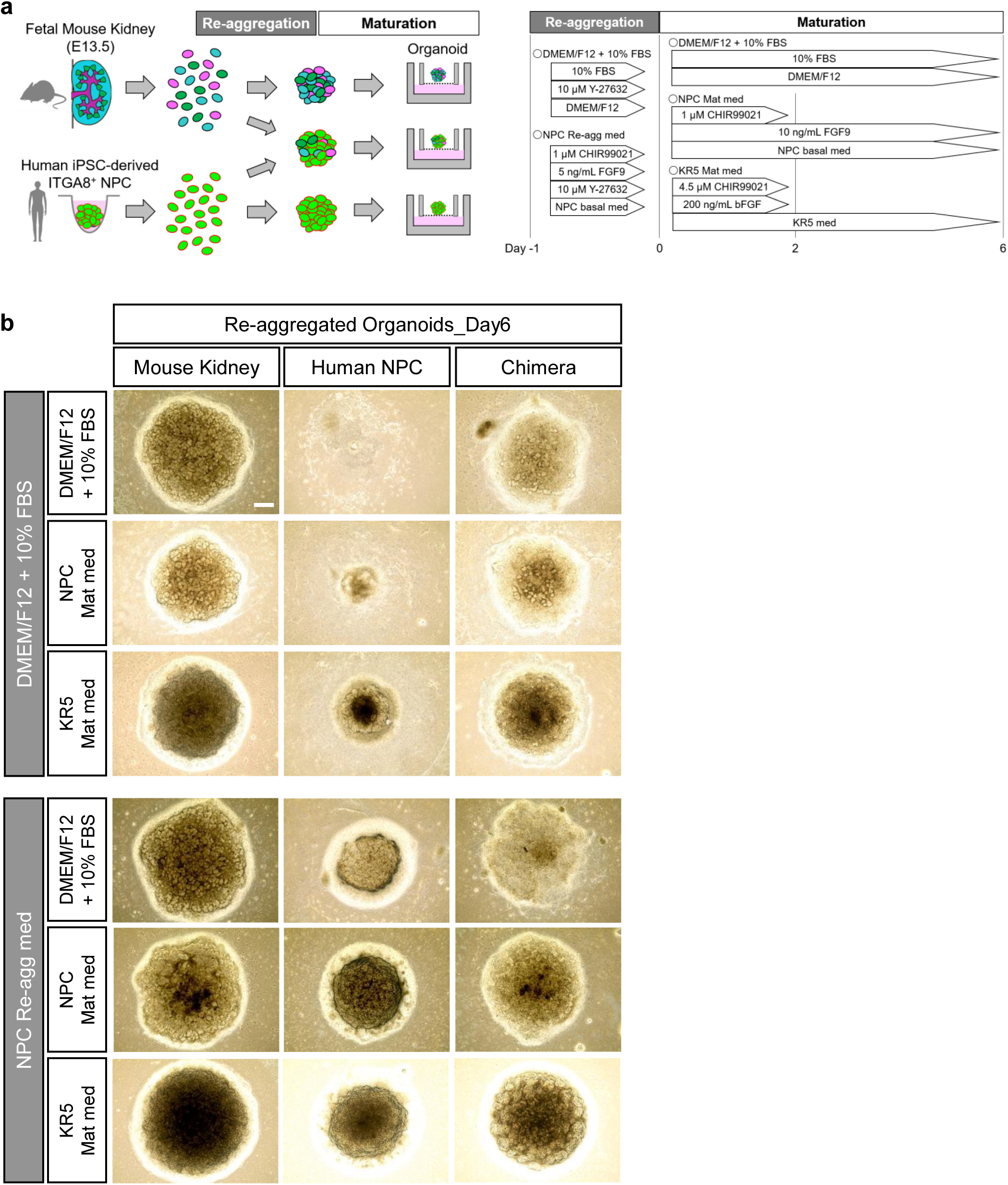
Screening of culture conditions using human-mouse chimeric renal organoids. a. Experimental scheme for the generation and examination of chimeric renal organoids under different culture conditions. b. Bright-field images of organoids formed under each culture condition. Scale bar represents 200 μm.

On the other hand, when utilizing NPC_Re-agg as the reaggregation medium, organoid formation was observed for all maturation conditions. Particularly, under the conditions employing the mature media NPC_Mat or KR5_Mat, the organoids exhibited a distinct three-dimensional structure (Fig. 1b). This dataset elucidated novel culture conditions, wherein three model renal organoids were formed: the mouse, human, and chimeric mouse-human model organoids. It was implied that both reaggregation and maturation media strongly influence renal organoid formation as well as chimeric renal organoid formation.

### Evaluation of heterogeneous chimeric renal organoids formed under different culture conditions

We conducted an immunostaining evaluation of six combinations of reaggregation and maturation media to investigate the conditions. Immunostaining was conducted on human-mouse chimeric organoids using representative markers of nephron segments, including WT1 (expressed in differentiating NPCs and glomeruli) and ECAD (specific to distal tubules), as well as the representative marker of ureteric buds (UBs), CK8. Regardless of the culture conditions, the formed human-mouse chimeric organoids exhibited WT1-positive glomerular structures, ECAD-positive distal tubule structures, and CK8-positive UB structures (Fig. 2a, b). No significant differences were observed in the CK8 positivity rate under any of the culture conditions (Fig. 2c). However, when NPC_Re-agg was used as the reaggregation medium, organoids exhibited a higher proportion of human cells and a greater density of kidney structures compared to organoids cultured in DMEM/F12 with 10% FBS (Fig. 2a, b). In the groups where KR5_Mat was used as the maturation medium, irrespective of the reaggregation medium used, numerous tubular structures were observed (Fig. 2a, b). Images of organoids costained for Ku80 and HuNu, human cell markers, demonstrated the formation of a “chimeric structure” including nephron structures that are part human and part mouse (Fig. 2a, b). These data suggest that the formed organoids are not mere mixtures of heterologous kidney cells but have common structures that develop in a coordinated manner.

**Fig. 2.**
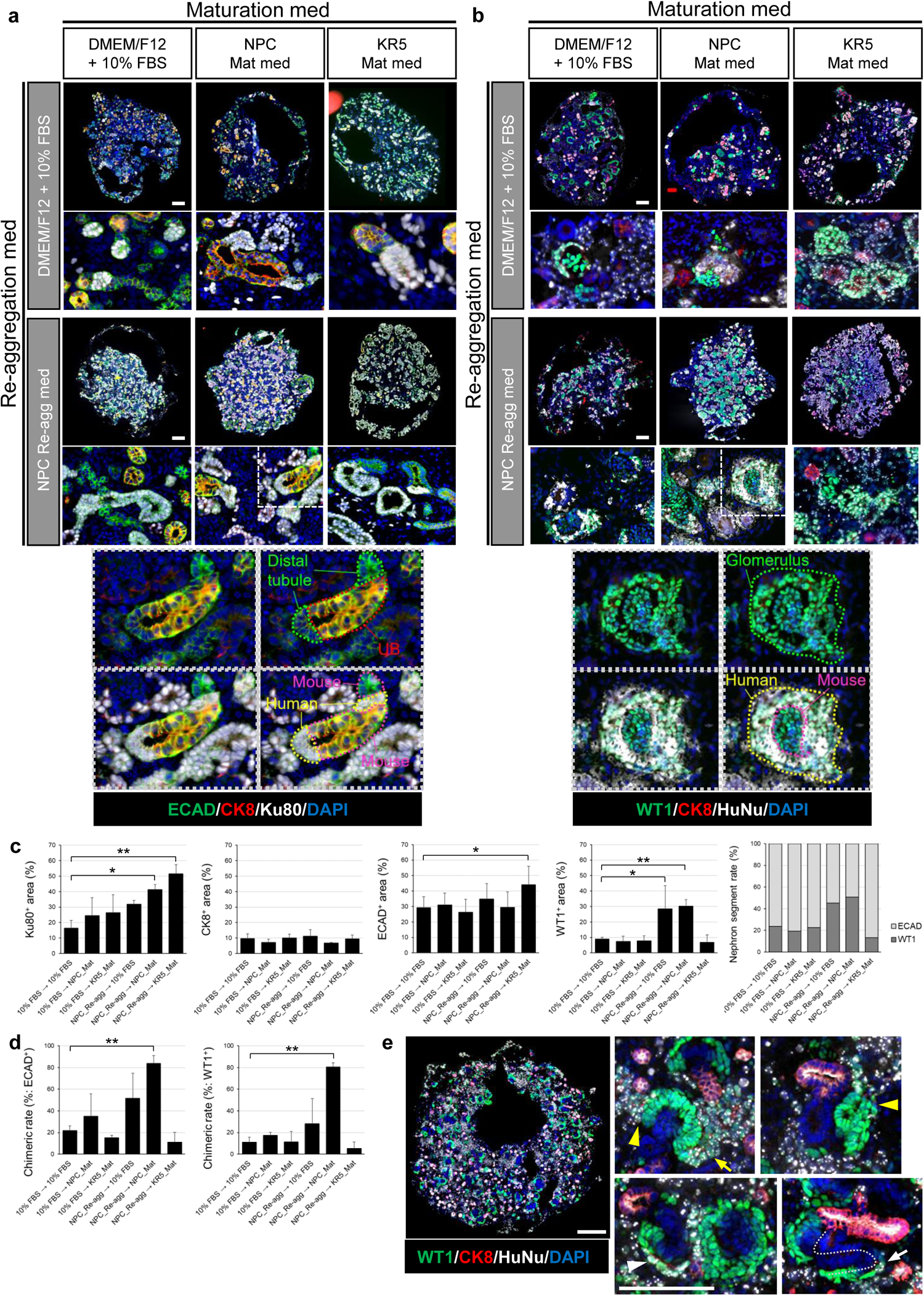
Evaluation of human-mouse chimeric renal organoids formed under different culture conditions. a. Immunostaining images of chimeric renal organoids at Day 6 that were stained using antibody against a distal tubule marker. Scale bar represents 200 μm. b. Immunostaining images of chimeric renal organoids at Day 6 that were stained using antibody against a glomerulus marker. Scale bar represents 200 μm. c. Cellular composition analysis based on immunostaining images (n= 3 independent experiments; mean ± s.d.; **P< 0.01; one-way ANOVA followed by the Tukey test). d. Quantitative analysis of chimera formation rate. The left graph indicates the chimera rate in distal tubule staining images, while the right graph indicates the chimera rate in glomerulus staining images (n= 3 independent experiments; mean ± s.d.; **P< 0.01; one-way ANOVA followed by the Tukey test). e. Immunostaining images of chimeric renal organoids at Day 2 of culture in the combination of NPC_Re-agg and NPC_Mat media. White arrow head indicates cap structure, yellow arrow heads indicate RVs, yellow arrow indicates comma-shaped body, and white arrow indicates S-shaped body. Scale bar represents 200 μm.

Chimeric renal organoids formation was demonstrated under all the investigated culture conditions. On the other hand, cell population analysis conducted on the 6th day of maturation culture revealed variations in the % area of human cells, ECAD-positive cells, and WT1-positive cells depending on the culture conditions (Fig. 2c). Particularly, using NPC_Re-agg as the reaggregation medium maintained the human cell content at a high level on the 6th day of maturation culture (Fig. 2c). This result emphasized the importance of the surrounding environment during reaggregation, and in contrast to the previously reported chimeric renal organoid reaggregation medium, the investigated medium in this study contained CHIR99021 and FGF9, suggesting that the activation of Wnt and FGF9 signals significantly contributed to human cell survival during chimeric renal organoid reaggregation.

Furthermore, using NPC_Re-agg as the reaggregation medium and NPC_Mat as the maturation medium, the highest nephron segment rate was achieved (Fig. 2c). On the other hand, even with NPC_Re-agg as the reaggregation medium, maturation with KR5_Mat resulted in a significant predominance of ECAD-positive cells (Fig. 2c), implying an influence on the differentiation and orientation of each nephron segment of NPCs.

Calculating the chimera formation rate of human-mouse chimeras, focusing on both glomeruli and distal tubules in the chimera evaluation, the combination of NPC_Re-agg and NPC_Mat significantly improved the chimera formation rate (Fig. 2d). Chimeric renal organoids cultured in the combination of NPC_Re-agg and NPC_Mat media exhibited chimeric organization, with human NPCs contributing to mouse cap structures at the early stage of mature culture, and chimeric structures characteristic of the renal vesicle (RV) (Fig. 2e and Supplementary Fig. 2), comma-shaped body, and S-shaped body stages (Fig. 2e).

The conditions utilizing NPC_Re-agg as the reaggregation medium and NPC_Mat as the maturation medium improved the formation of human, mouse, and human-mouse chimeric organoids, compared to the previous report[20]. Additionally, the composition of the medium was shown to influence the ratio of components in the chimera.

### Evaluation of fetal pig kidney development and formation of fetal pig kidney organoids

Next, we evaluated whether the selected culture conditions were suitable for forming organoids using fetal pig kidney cells. Initially, fetal pig kidneys were selectively collected through microscopic surgery of E30 porcine fetuses immediately after their removal from the uterus. The size and morphology of the harvested E30 porcine kidneys were similar to those of conventionally used E13.5 mouse kidneys. Multiple cap structures were observed beneath the renal capsule in the porcine E30 kidneys (Fig. 3a). Morphological analysis indicated that while the E13.5 mouse kidneys used in this study showed nephron development up to the RV stage, the E30 porcine kidneys exhibited structures progressing up to the comma-shaped and S-shaped body stages, albeit with some variation among individual porcine fetuses (Fig. 3a). These data suggest a slightly more advanced stage of development in E30 porcine kidneys compared to E13.5 mouse kidneys used in this study (Fig. 3a).

**Fig. 3.**
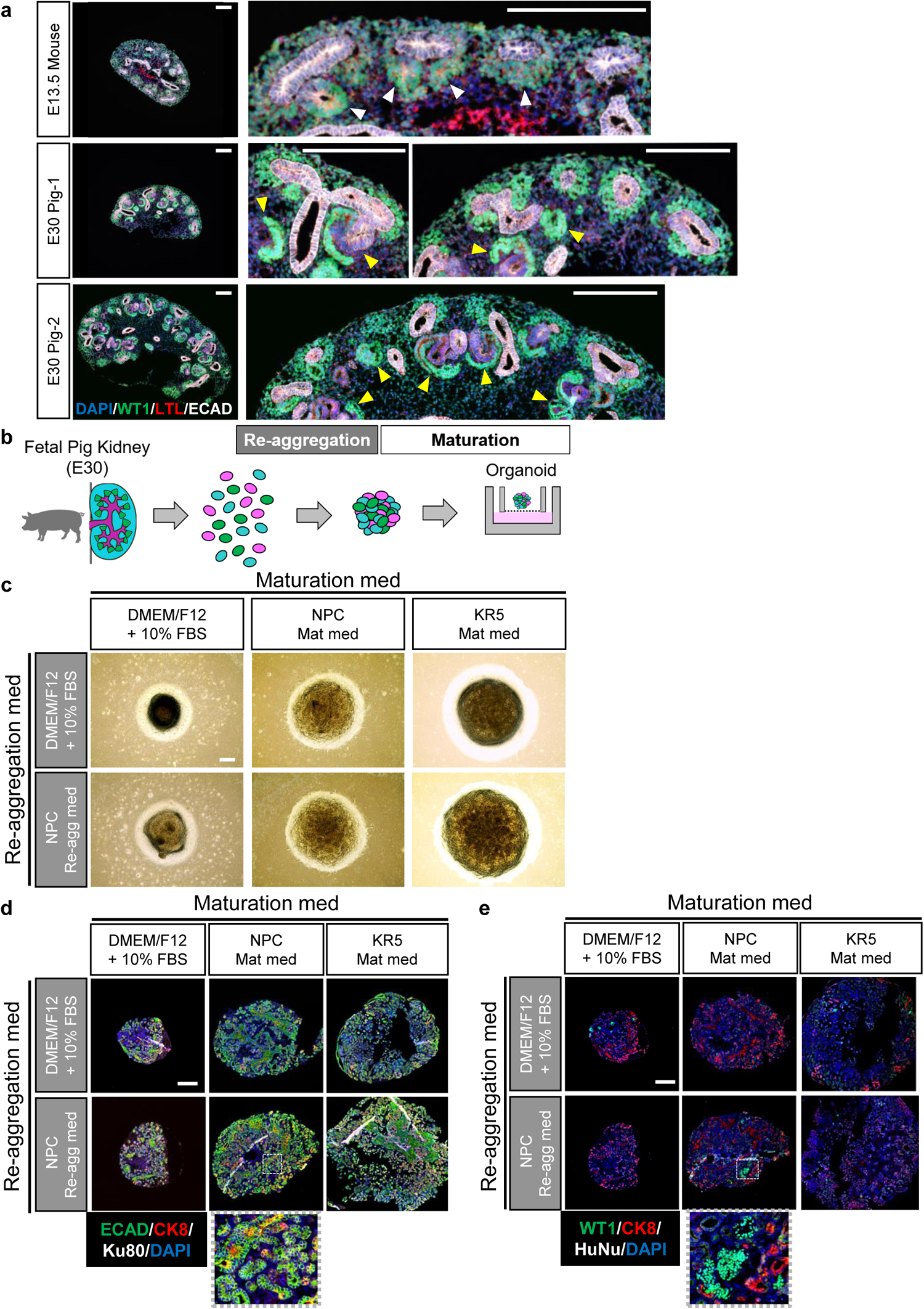
Generation and evaluation of fetal pig kidney organoids. a. Histological comparative analysis of E13.5 mouse kidney and E30 pig kidney. White arrow heads indicate RVs and yellow arrow heads indicate comma-shaped bodies or S-shaped bodies. Scale bar represents 200 μm. b. Experimental scheme of organoid formation using fetal pig kidneys. c. Bright-field images of pig kidney organoids formed under different culture conditions. Scale bar represents 200 μm. d. Immunostaining images of pig kidney organoids at Day 6 that were stained by antibody against a distal tubule marker. Scale bar represents 200 μm. e. Immunostaining images of pig kidney organoids at Day 6 that were stained by antibody against a glomerular marker. Scale bar represents 200 μm.

Similar to the study in mice, single-cell dissociation was performed on porcine E30 kidneys, and reaggregation and maturation media were compared. Organoids were formed under all culture conditions (Fig. 3b, c). However, the organoid sizes varied with each culture condition. Regardless of the reaggregation medium used, larger porcine fetal organoids were formed when the maturation medium was either NPC_Mat or KR5_Mat, compared to the conventional medium (Fig. 3b, c).

Immunostaining analysis conducted on the 6th day of maturation revealed differences in the sizes of porcine fetal kidney organoids based on the culture medium (Fig. 3d, e). However, all porcine fetal kidney organoids were confirmed to possess kidney structures positive for WT1, ECAD, and CK8 (Fig. 3d, e). Moreover, irrespective of the culture condition, ECAD-positive structures predominated over WT1-positive ones, and a bias towards the distal tubules was observed in nephron segments (Fig. 3d, e). The combination of reaggregation medium NPC_Re-agg and maturation medium NPC_Mat yielded the highest number of WT1-positive cells (Fig. 3d, e).

### Formation and evaluation of human-pig chimeric renal organoids

Based on the results of the porcine fetal kidney organoid culture conditions (Fig. 3b-e) and the evaluation of human-mouse chimeric renal organoids (Fig. 1b, 2a-e), we selected the combination of reaggregation medium NPC_Re-agg and maturation medium NPC_Mat for initiating the cultivation of human-porcine chimeric renal organoids (Fig. 4a). We proceeded with the cultivation of human-porcine chimeric renal organoids, varying the human-porcine cell mixture ratios to observe their impact on chimeric organoid formation. The cell resources included E30 porcine fetal kidneys and human iPSC-derived NPCs, the latter having been further enriched using magnetic-activated cell sorting (MACS). We confirmed that these ITGA8-positive NPCs derived from human iPSCs exhibited the potency to differentiate and mature into nephrons (Fig. 4b).

**Fig. 4.**
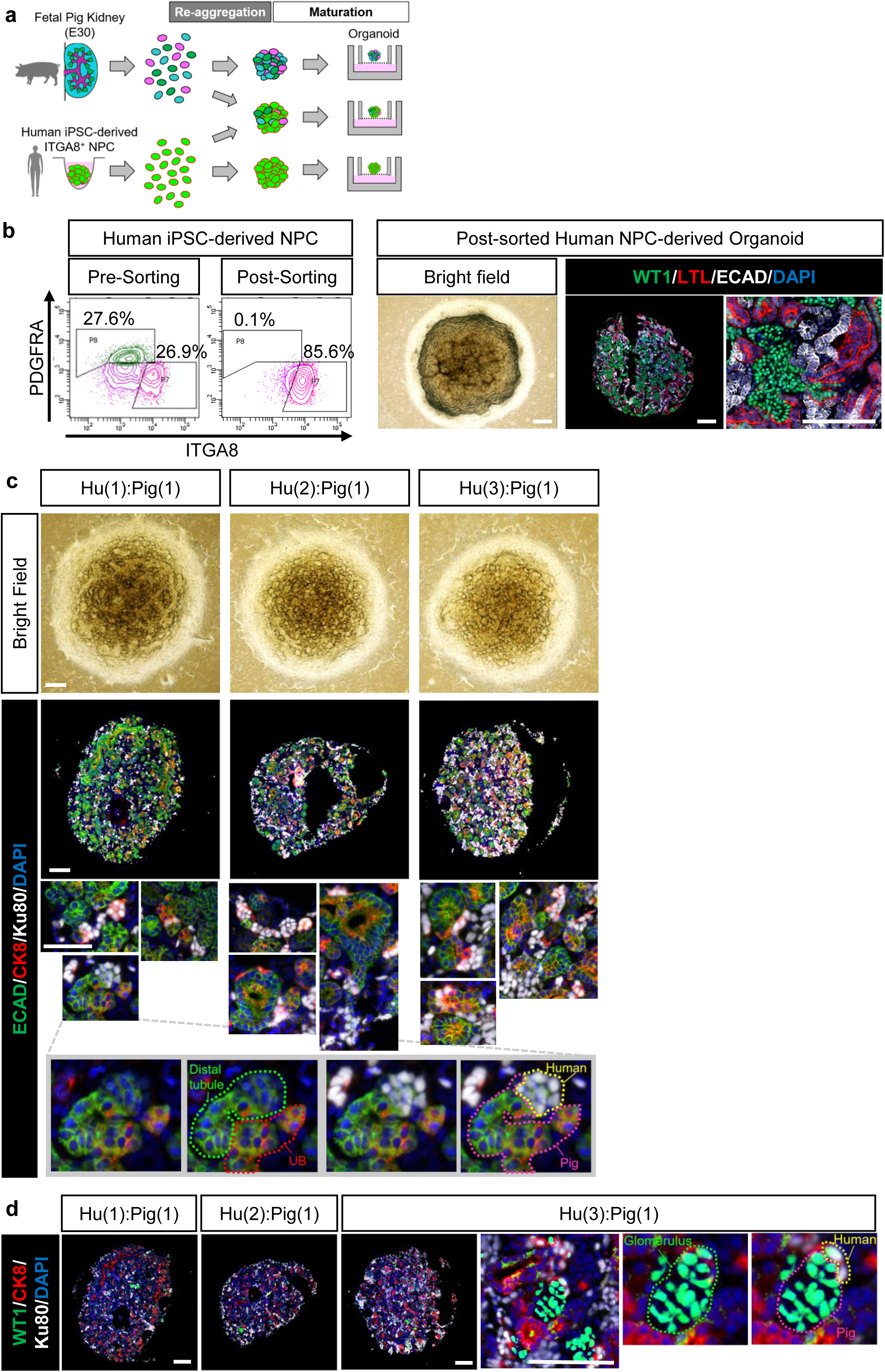
Generation and evaluation of human-pig chimeric renal organoids. a. Experimental scheme for generating human-pig chimeric renal organoids using fetal pig kidneys and human iPSC-derived NPCs. b. Quality assessment of human NPCs by flow cytometry and evaluation of differentiation potency of sorted NPCs. Scale bar represents 200 μm. c. Bright-field images and immunostaining images of chimeric renal organoids formed at different human-pig mixing ratios and stained by antibody against a distal tubule marker. Scale bar represents 200 μm. d. Bright-field images and immunostaining images of chimeric renal organoids formed at different human-pig mixing ratios and stained by antibody against a glomerular marker. Scale bar represents 200 μm.

By utilizing these two cell resources and cultivating them under the selected culture conditions, we confirmed the formation of human-porcine chimeric renal organoids (Fig. 4c). Immunostaining using a distal tubule marker revealed the formation of chimeric structures in multiple locations within the human-porcine chimeric renal organoids (Fig. 4c).

Immunostaining analysis using a glomerular marker showed that glomeruli were present in very few numbers within the human-porcine chimeric renal organoids (Fig. 4d). Similar findings were observed in porcine fetal kidney organoids (Fig. 3d, e), suggesting that the origin of this phenomenon was likely the porcine fetal kidney rather than the human NPC component. However, the few glomerular structures identified had chimeric structure, indicating that human NPCs demonstrated chimera-forming ability within porcine glomeruli (Fig. 4d).

### Utilization of selected culture conditions for generation of humanized xenogeneic kidneys

In our previous study, we confirmed that human NPCs can mature and connect with mouse UBs, reaching the RV stage when cultured in MEMα medium supplemented with FBS [16]. However, the engraftment efficiency of human NPCs was low, and the formation of chimeric structures was limited. Therefore, in this study, we investigated whether the engraftment and chimera formation abilities of human NPCs could be enhanced by culturing them in selected conditions using xenogeneic embryonic kidneys after injection (Fig. 5a).

**Fig. 5.**
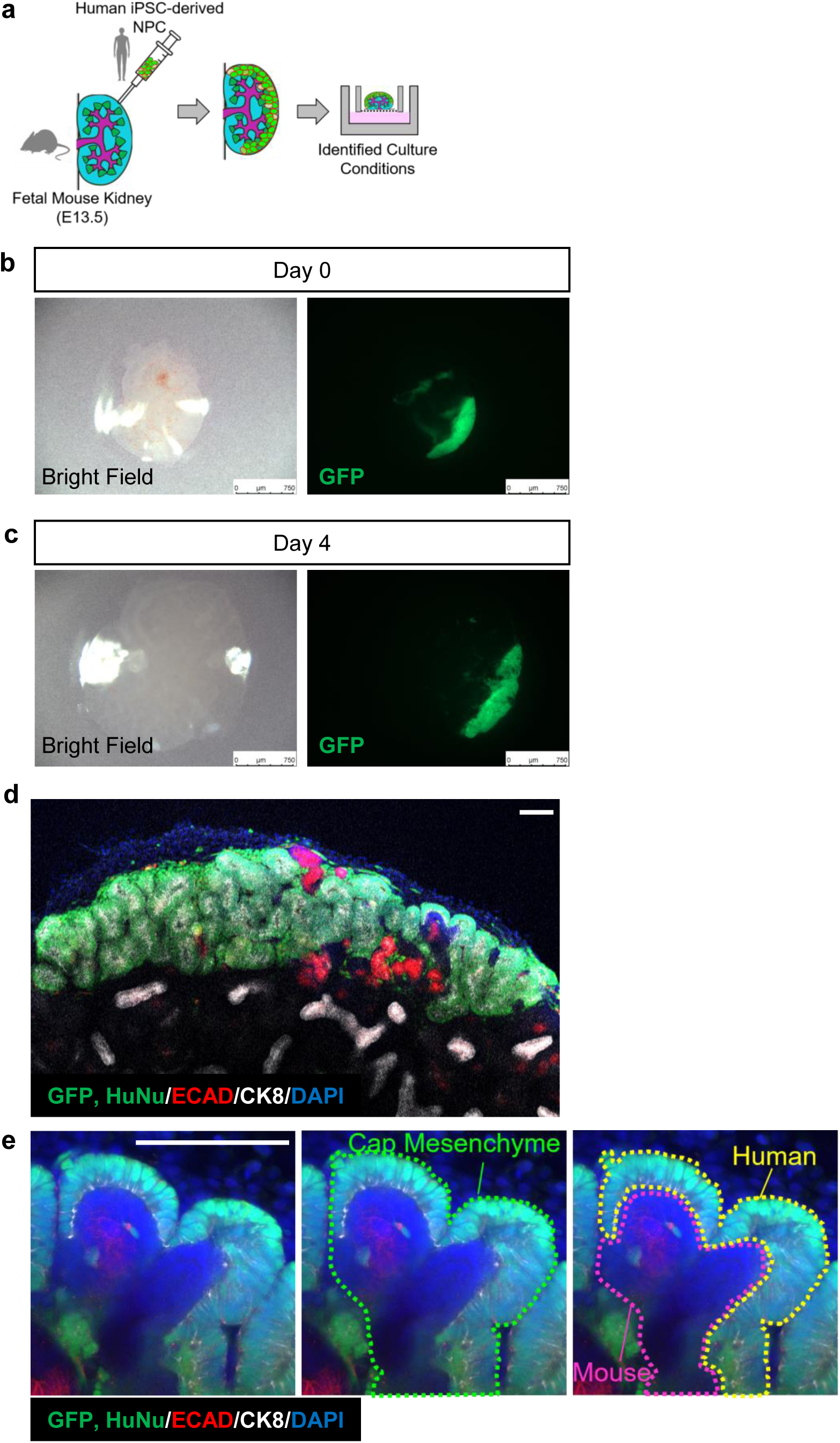
Evaluation of human NPC-injected fetal mouse kidney using specific culture conditions. a. Experimental scheme for the generation and *in vitro* culture of human-mouse chimeric kidneys. b. Fluorescence stereomicroscope images of fetal mouse kidney immediately after injection of human NPC. Scale bar represents 750 μm. c. Fluorescence stereomicroscope images of human NPC-injected fetal mouse kidney cultured for 4 days under the identified culture conditions. Scale bar represents 750 μm. d, e. Immunostaining images of human NPC-injected fetal mouse kidney cultured for 4 days under the identified culture conditions. Scale bar represents 100 μm.

EGFP-labeled human iPSC-derived NPCs were dissociated into single cells and injected under the renal capsule of E13.5 mouse embryos, which were then cultured at the air-liquid interface in NPC_Re-Agg medium for four days. Microscopic examination of the injected fetal mouse kidneys immediately after injection revealed that the morphology of the recipient kidneys remained intact, and the amount of EGFP-labeled human NPCs injected beneath the renal capsule surface was sufficient (Fig. 5b). By the fourth day of culture, the injected fetal mouse kidneys exhibited increased size and flattening, while the human NPCs injected initially remained abundantly present beneath a wide area of the renal capsule surface (Fig. 5c).

Immunostaining analysis revealed a thick and layered engraftment of human cells beneath the mouse embryonic renal capsule, forming a three-dimensional structure (Fig. 5d and Supplementary Movie 1). Furthermore, high-magnification images clearly demonstrated the engraftment of human cells at the tip of the mouse UB, strongly indicating the formation of chimeric cap mesenchyme (Fig. 5e and Supplementary Movie 2). These results demonstrate that the selectively chosen culture conditions contribute to the survival and chimera formation ability of human NPCs injected into mouse fetal kidneys beneath the subrenal capsule, thus accelerating the pace of future research on the generation of humanized xenogeneic chimera kidneys.

## Discussion

This study established an optimized chimeric renal organoid culture system by refining the existing culture conditions for human-mouse chimeric renal organoids, dividing them into reaggregation stage and maturation stage culture conditions, and optimizing the medium composition. This optimized culture system allows both the animal and human cellular components to survive long-term and facilitates more efficient interspecies chimera formation. Furthermore, the identified optimized culture conditions were found to be suitable for the formation of porcine fetal kidney organoids. Ultimately, we achieved the first successful creation of human-porcine chimeric renal organoids. The ability of human NPCs to form chimeric renal organoids with porcine cells strongly supports the potential of utilizing the developmental environment of porcine fetus, such as through fetal organ complementation, to enable the development of human renal precursor cells. In addition, we discovered that using selected conditions to culture fetal mouse kidneys, into which human NPCs were subcapsularly injected, improved the survival of human cells and enhanced chimera formation. These findings hold significant promise for future research on the production of human-interspecies chimeric kidneys for clinical purposes.

In order to improve cell survival and chimera formation rates in human-mouse chimeric renal organoids and achieve the formation of human-pig chimeric renal organoids, we initially explored conditions for the stable organoid maturation of the constituent cells, human NPCs and fetal mouse (or pig) kidney cells. Through the investigation of conditions, we identified culturing conditions where cells from different animal species can survive long-term within the chimeric renal organoids. Consequently, it is believed that the opportunities for interspecies cell-cell interactions increased, leading to a higher chimera formation rate. These findings suggest that the optimization of culture conditions can surpass the barrier mechanisms that suppress interspecies chimera formation[22], indicating a significant developmental insight.

In particular, a significant improvement in the human cell ratio within chimera organoids was observed when NPC_Re-agg was used as a re-aggregation culture medium (Fig. 2c). FGF9, which is added to NPC_Re-agg, has been reported to contribute to the maintenance and proliferation of human NPCs[30]. Therefore, it is believed that the FGF9 signaling pathway influenced the enhancement of the human cell ratio in chimera organoids. Recently, Li et al. successfully induced the differentiation of renal organoids from naïve-like embryonic stem cells (ESCs) derived from pigs[31]. Although our study differs from theirs in terms of the original cellular resources used, they also employed FGF9 and Wnt signal activators in the process of porcine renal organoid generation. Considering that their porcine ESC-derived renal organoids were produced by appropriately controlling both signals, which is analogous to the human kidney development process[32–34], it is suggested that FGF9 and Wnt signals are also important factors in pig kidney development. Thus, the observed commonalities between human and pig kidney development processes are believed to be one of the contributing factors facilitating the coexistence and chimera formation of human NPCs and pig fetal kidney cells in our study.

On the other hand, when KR5_Mat was used as the maturation medium, although the survival rate of human NPCs and fetal mouse kidney cells was maintained, the chimera formation rate actually decreased (Fig. 2d). KR5_Mat provides a strong Wnt signal in the early stages of maturation culture, and it has been reported that excessive Wnt signal not only accelerates nephron development but also biases differentiation towards the distal tubules[35–37]. Furthermore, Chen et al. demonstrated through experiments involving the injection of embryonic mouse NPCs into cap mesenchyme of mouse fetal kidneys at different developmental stages, the developmental timing difference significantly influences the survival and differentiation of injected NPCs[38].

Thus, it is postulated that the strong action of Wnt signal disrupted the balance of self-renewal and differentiation of human NPCs, leading to an increased population of epithelialized or differentiated NPCs, as well as causing a mismatch in developmental stages between human and mouse. Consequently, this perturbation is believed to have resulted in a reduction of opportunities for cell-cell interactions in the appropriate cellular state and mistiming of chimera formation. In cells cultured in NPC_Mat medium, which maintains a relatively low Wnt signal condition, no bias in differentiation was observed, and efficient chimera formation was achieved (Fig. 2d). This suggests that, not only cell survival rates but also the regulation of differentiation direction, as well as the synchronization of developmental stages between mice and humans as previously discussed[39,40], may be important factors for chimera formation. Further research, considering differences among animal species, is anticipated to shed more light on this matter.

Differences in the composition of nephron segments were observed between porcine and murine fetal kidney organoids. In this study, E30 pig and E13.5 mouse kidneys were used as cellular resources, considering potential limitations in the accuracy of pregnancy detection method and inter-individual variations. Nonetheless, at least in the case of E30 porcine kidney, a greater abundance of late-stage nephron structures was observed. Furthermore, in E13.5 mouse kidney organoids, an increase in the proportion of distal tubules dependent on Wnt signaling during culture was observed (data not shown), while in fetal pig kidney organoids, the composition of cell types remained relatively stable regardless of the strength of Wnt signaling (Fig. 3d, e). Considering the decrease in NPCs and the increase in tubular structures as nephron development progresses, the observed differences in the nephron segment composition in this study cannot be entirely disregarded as solely due to the differences in animal species. It is believed that, at the very least, variations in the developmental origins have influenced these differences.

The chimeric renal structures observed in this study were mosaic chimeras, where some of the renal structures were replaced by the host animal components. This is likely attributed to the fact that the host animal NPCs were not excluded in this experimental system. The future development of a system that selectively excludes host pig NPCs, such as Six2-DTA systems[16], is expected to further enhance the chimera formation rates and achieve complete humanization of NPCs in the porcine kidney.

Given that NPCs are known to exist beneath the renal capsule during the fetal period, we have endeavored to generate humanized kidneys by injecting human NPCs beneath the capsule of xenogeneic fetal kidneys. In this study, we have successfully identified culture conditions that improve the survival and chimera formation of not only organoids but also human NPCs injected beneath the renal capsule. This finding could serve as a pivotal milestone towards the generation of xenogeneic chimera kidneys in the future.

In summary, this study provides foundational technology that can significantly accelerate future research on the creation of human-porcine chimeric kidneys. It also holds promise for utilization as an evaluation technique to ensure the quality of human NPCs especially for clinical application.

## Methods

### Human iPSC culture

The human iPSC line, iPSC-S09, and EGFP-labeled human iPSC line, 317-12-Ff[41], were used in this study. The iPSC-S09 line was established by Sumitomo Pharma Co., Ltd. from peripheral blood cells using Sendai virus vectors. The cells were maintained on iMatrix-511 (Nippi. Inc., Osaka, Japan) in StemFit medium (Ajinomoto Co., Inc, Tokyo, Japan), and cultured in a humidified atmosphere of 5% CO_2_ at 37°C. The cell culture medium was changed every 1-2 days, and passaged every 7 days by treatment with TrypLE Select Enzyme (Thermo Fisher Scientific Inc., MA, USA).

The experimental protocols using clinical grade human iPSCs were approved by the Research Ethics Committee of Sumitomo Pharma Co., Ltd., Japan.

### Induction of NPCs from human iPSCs

NPCs were induced from human iPSCs according to previously established methods[28,42] with slight modification. Briefly, on the day of induction (day 0), 10,000 iPSCs per well were seeded in a PrimeSurface 96 Slit-Well plate (Sumitomo Bakelite Co., Ltd., Tokyo, Japan) containing basal medium for human NPCs (NPC basal medium)[28,42], which was supplemented with 3 ng/mL human activin A (R&D Systems, Inc., Minneapolis, MN, USA), 1 ng/mL human bone morphogenetic protein 4 (Bmp4) (R&D Systems, Inc.), 20 ng/mL human basic fibroblast growth factor (b-FGF) (R&D Systems, Inc.), and 10 μM Y27632 (FUJIFILM Wako Pure Chemical Corporation, Osaka, Japan). After 24 h of incubation (day 1), the culture medium was changed to the NPC basal medium with 10 μM CHIR99021 (FUJIFILM Wako Pure Chemical Corporation). Every other day thereafter (days 3 and 5), half of the medium was replaced with NPC basal medium containing 10 μM CHIR99021 (FUJIFILM Wako Pure Chemical Corporation) and 10 μM Y27632 (FUJIFILM Wako Pure Chemical Corporation). On day 7, the medium was switched to the NPC basal medium, which was supplemented with 10 ng/mL human activin A (R&D Systems, Inc.), 5 ng/mL human Bmp4 (R&D Systems, Inc.), 3 μM CHIR99021, 0.1 μM retinoic acid (Sigma-Aldrich, MO, USA), and 10 μM Y27632 (FUJIFILM Wako Pure Chemical Corporation). On day 10, the medium was changed to the NPC basal medium, which was supplemented with 1 μM CHIR99021 (FUJIFILM Wako Pure Chemical Corporation), 5 ng/mL fibroblast growth factor-9 (FGF-9) (Abcam plc., Cambridge, United Kingdom), and 10 μM Y27632 (FUJIFILM Wako Pure Chemical Corporation). On day 13, the spheres were collected for experimental use.

The NPCs used for injection into the fetal mouse kidney were induced following a previously reported method[43].

### Cell sorting

For the selection of integrin subunit alpha 8 (ITGA8)-positive NPCs, anti-Biotin MicroBeads UltraPure (Miltenyi Biotec B.V. & Co. KG, Bergisch Gladbach, Germany), MS Columns (Miltenyi Biotec B.V. & Co. KG), and a MiniMACS Separator (Miltenyi Biotec B.V. & Co. KG) were used according to the manufacturer’s protocols.

### Research animals

The experiments were conducted in accordance with the National Institutes of Health Guide for the Care and Use of Laboratory Animals. Every effort was made to minimize animal suffering. Pregnant female C57BL/6NCrSlc mice on embryonic day 13.5 were purchased from Japan SLC, Inc. (Shizuoka, Japan). Pregnant female Microminipigs on embryonic day 30 were purchased from Fuji Micra, Inc. (Shizuoka, Japan).

### Single cell extraction from fetal kidneys

Single cells from mouse and pig fetal kidneys were obtained following a previously reported method[20,25,44]. Fetal kidneys were harvested from the decapitated fetuses under an operating microscope and collected in 1.5 mL tubes containing a minimum essential medium (MEMa; Thermo Fisher Scientific, Inc.). The tube was then centrifuged at 700 g for 3 min, the supernatant was removed, and 1 mL of accutase (AT104, Innovative Cell Technologies, Inc. CA, USA) was dispensed. The sample was vortexed and incubated at 37°C for 5 min (repeated twice), and further manual pipetting and incubation was performed again for 5 min. The cell suspension was then centrifuged at 300 g for 5 min to remove the subsequent accutase supernatant. The pellet was resuspended in 1 mL of PBS (Thermo Fischer Scientific, Inc.) with 10 μM Y27632 (FUJIFILM Wako Pure Chemical Corporation). In addition, the cells were passed through a 40 μm cell strainer (Corning Incorporated, NY, USA) to remove clumps of cells and obtain a single-cell suspension of mouse or pig cells.

### Chimeric renal organoids culture

In accordance with the methods described above, ITGA8-positive human NPCs, as well as dissociated fetal mouse kidney cells or fetal pig kidney cells, were prepared.

The cell density for all samples was adjusted to 1 × 10^6^ cells/mL in DMEM/F12 medium (Thermo Fisher Scientific Inc.) with 10% FBS and 10 μM Y27632 (FUJIFILM Wako Pure Chemical Corporation) or NPC basal medium with 1 μM CHIR99021 (FUJIFILM Wako Pure Chemical Corporation), 5 ng/mL FGF-9 (Abcam plc.), and 10 μM Y27632 (FUJIFILM Wako Pure Chemical Corporation), referred to as “NPC_Re-agg”.

The human-mouse mixture was prepared in a ratio of 1:3, and the human-pig mixtures were prepared at ratios of 1:1, 2:1, or 3:1, respectively. The mixtures were seeded into a V-bottomed 96-well low-binding plate (Sumitomo Bakelite Co., Ltd.) to achieve a total of 2 × 10^5^ cells/well containing 200 µL of culture medium. The 96-well plate was centrifuged at 1,000 rpm for 4 min and cultured overnight in a 5% CO_2_ incubator at 37°C.

The formed mixture aggregates were transferred to the air-liquid interface and cultured for a maximum of 6 days to promote maturation. As the maturation medium, we used DMEM/F12 medium (Thermo Fisher Scientific Inc.) with 10% FBS, NPC basal medium with 1 μM CHIR99021 (FUJIFILM Wako Pure Chemical Corporation) and 10 ng/mL FGF-9 (Abcam plc.), referred to as “NPC_Mat”, or KR5 medium[26,27] with 4.5 μM CHIR99021 (FUJIFILM Wako Pure Chemical Corporation) and 200 ng/mL bFGF (R&D Systems, Inc.), referred to as “KR5_Mat”. After 48 hours of maturation culture, NPC_Mat medium without CHIR99021 and KR5_Mat medium without CHIR99021 and bFGF were prepared and exchanged.

The same method as described above was employed even in the case of non-chimeric renal organoid cultures.

### Injection of human NPCs into the subrenal capsule of fetal mice

Human iPSC-derived NPCs, without cell sorting, were injected under the capsule of the embryonic kidney according to a previously reported method[17]. In detail, C57BL/6NCrSlc mice were euthanized by cervical dislocation at 13.5 days of gestation. The fetuses were then removed with the uterus and placed in a 10-cm dish containing Hank’s Balanced Salt Solution. Thereafter, the fetuses were separated from the uterus and euthanized via decapitation. First, a cut was made along the spinal cord from the base of the hindlimb of the fetus, and a similar cut was made on the other side to take out the vertebral column. After the kidneys on both sides of the fetus were visible, the fetus was fixed with micro-tweezers. Subsequently, progenitor cells were placed in glass needles. Afterward, the glass needles were inserted into the renal capsule through the renal hilus under a fluorescent stereomicroscope (Leica M205FA, Leica Microsystems GmbH, Wetzlar, Germany) and the human NPCs were injected under the renal capsule by mouth pipetting. The injected kidneys were detached from the fetus and then transferred to the air-liquid interface for 4 days of *in vitro* culture. As the organ culture medium, we used NPC basal medium with 1 μM CHIR99021, 5 ng/mL FGF-9, and 10 μM Y27632, referred to as “NPC_Re-agg”. After 24 hours of culture, NPC_Re-agg medium without Y27632 was prepared and exchanged.

### Frozen section immunostaining

*In vitro* cultured organoids, fetal mouse kidneys, and fetal pig kidneys were fixed with 4% paraformaldehyde (FUJIFILM Wako Pure Chemical Corporation) and sectioned with a cryostat (Leica Microsystems GmbH) to prepare frozen sections. Frozen sections were treated with antigen retrieval reagent (RM102-H, LSI Medience Corporation, Tokyo, Japan) at 121°C for 5 min. The primary antibodies used were as follows: anti-Wilms’ tumor-1 (WT1; 1:400; ab89901, Abcam plc. / 1:100; sc-7385, Santa Cruz Biotechnology Inc., TX, USA), anti-cytokeratin 8 (CK8 (TROMA-I); 1:100, DSHB, IA, USA), anti-human nuclei (HuNu; 1:100; MAB4383, Merck Millipore, MA, USA), anti-Ku80 (1:100; 2180S, Cell Signaling Technology, Inc., MA, USA), anti-lotus tetragonolobus lectin (LTL; 1:500; B-1325, Vector Laboratories, Inc., CA, USA), and anti-E-cadherin-1 (ECAD; 1:500; 610181, BD Biosciences, CA, USA). Nuclear counterstaining was performed with DAPI (Nacalai Tesque, Inc., Kyoto, Japan). Stained sections were analyzed with the imaging cytometer, IN Cell Analyzer 6000 (Cytiva, Tokyo, Japan). Image analyses were performed with the imaging software, IN Cell Developer Toolbox (Cytiva).

### Whole-mount immunostaining

Cultured fetal mouse kidneys injected with human NPCs were fixed with 4% paraformaldehyde in PBS for 15 min at 4°C and washed three times with PBS. Samples were blocked using 1% donkey serum, 0.2% skimmed milk, and 0.3% Triton X/PBS for 1 h at room temperature, and incubated overnight at 4°C with primary antibodies. The primary antibodies used were as follows: anti-cytokeratin 8 (CK8 (TROMA-I); 1:100, DSHB), anti-human nuclei (HuNu; 1:100; MAB4383, Merck Millipore), and anti-E-cadherin-1 (ECAD; 1:500; 610181, BD Biosciences). After washing three times with PBS, the samples were incubated with secondary antibodies for 1 h at room temperature. Samples were mounted with ProLong Gold Antifade Mountant containing DAPI (P36931, Thermo Fisher Scientific Inc.). Each sample was examined under an LSM880 confocal laser scanning microscope (Carl Zeiss AG, Baden-Württemberg, Germany).

### Measurement of the chimeric rate

Immunostained sections of frozen organoids were captured using a fluorescence microscope. Chimeric structures were defined as renal structures with ECAD-positive cells containing Ku80-positive cells and renal structures with WT1-positive cells containing HuNu-positive cells. The number of chimeric structures and the total number of renal structures were quantified. The chimera formation rate was calculated by dividing the number of chimeric structures by the total number of renal structures. For chimera rate calculation, three sections from the central slice of the organoid specimen and two additional sections located 50 µm apart from anterior and posterior directions were used. Three independent experiments were conducted.

### Area-based cell composition analysis

Immunostained sections of frozen organoids were captured using a fluorescence microscope. Using the IN Cell Developer Toolbox (Cytiva), the fluorescence intensity of the organoid slice images was used to calculate the positive areas of ECAD, WT1, CK8, Ku80, and DAPI. The positive area of each marker was divided by the DAPI-positive area to calculate the proportion of each constituent cell type within the organoid. For cell composition analysis, three sections from the central slice of the organoid specimen and two additional sections located 50 µm apart from anterior and posterior directions were used. Three independent experiments were conducted.

### Flow cytometry

Biotinylated anti-ITGA8 (R&D Systems, Inc.), allophycocyanin-labeled streptavidin (BioLegend, Inc., CA, USA), and phycoerythrin-labeled anti-platelet-derived growth factor receptor A (PDGFRA) (BioLegend, Inc.) were used for cell staining. Data were acquired and analyzed using BD FACSAria IIu (BD Biosciences) and BD FACSDiva software (BD Biosciences), respectively.

### Statistical analysis

Statistical analyses were performed with R version 3.6.0 (The R Foundation for Statistical Computing). A two-tailed Student t-test was carried out for two-group comparisons, and one-way analysis of variance (ANOVA) followed by a Tukey test was performed for multiple-group comparisons.

## Acknowledgements

We thank Yuna Yakura and Mizuki Otake for experimental and technical assistance.

This work was supported by the Japan Agency for Medical Research and Development (AMED; grant no. 22bm1223003h0001 and 23bm1123036h0001) and JST FOREST Program (grant no. JPMJFR2011).

## Author contributions

K.F., S.Y., K.S., K.M., and T.K designed the study. K.F., S.Y., K.S., K.M., S.K., and T.K. carried out experiments and analyzed the data. K.F. wrote the manuscript. S.Y., K.S., K.M., T.K, A.I., M.I., E.K., and T.Y. interpreted the data and revised the manuscript. A.I., M.I., E.K., and T.Y. supervised the project. All authors have approved the final version of the manuscript.

## Data availability statement

All reasonable requests will be promptly reviewed by the corresponding authors to determine whether the request is subject to any intellectual property or confidentiality obligations.

---

**Supplementary Fig. 1.**
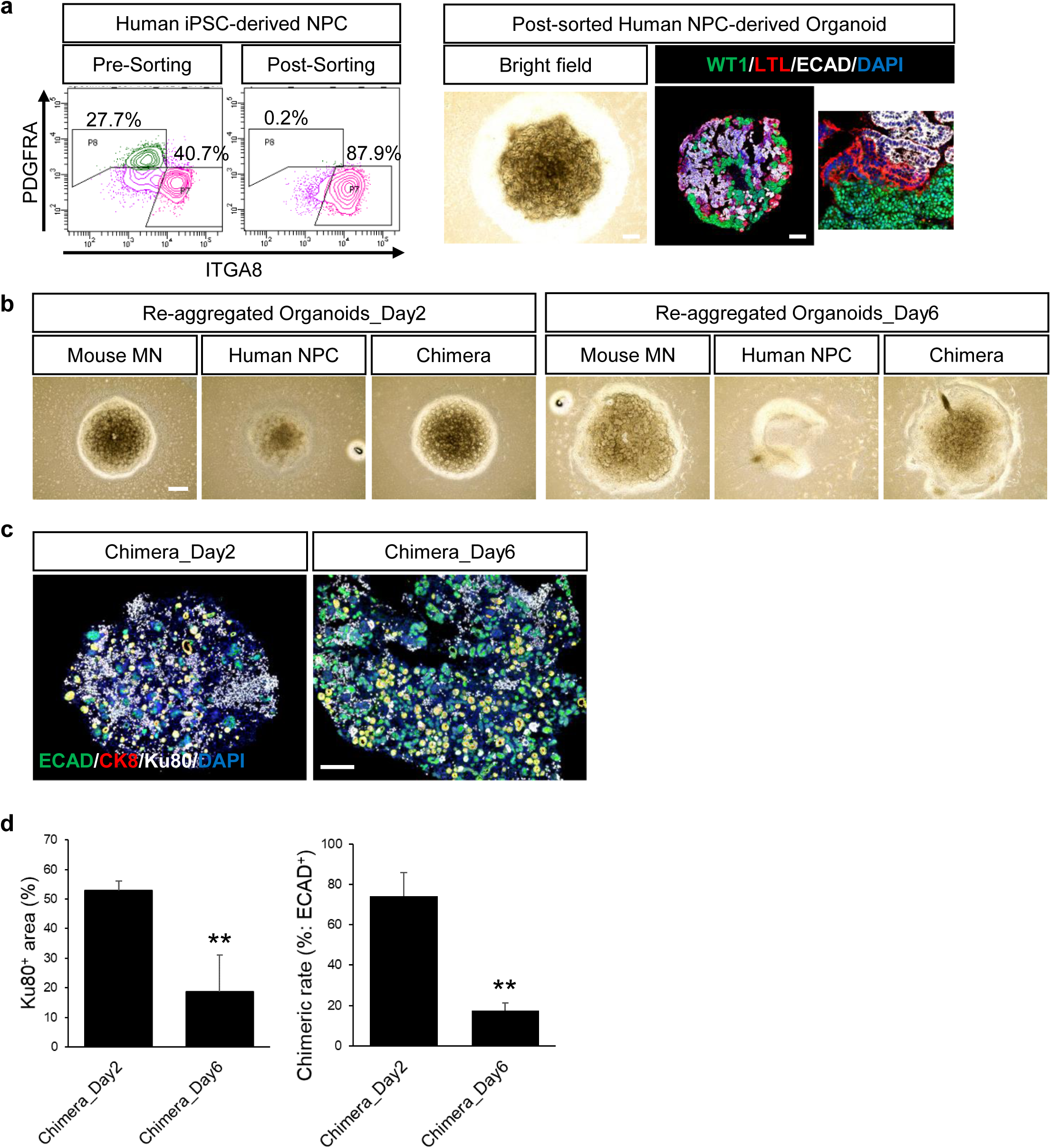
Evaluation of human-mouse chimeric renal organoids generated using existing protocols. a. Quality assessment of human NPCs used for human-mouse chimeric renal organoid experiments. Scale bar represents 200 μm. b. Bright-field images of human-mouse chimeric renal organoids, fetal mouse kidney organoids, and human NPC organoids generated using existing protocols. Scale bar represents 200 μm. c. Immunostaining images of human-mouse chimeric renal organoids, fetal mouse kidney organoids, and human NPC organoids generated using existing protocols. Scale bar represents 200 μm. d. Cell composition analysis and chimera formation analysis based on immunostaining of human-mouse chimeric renal organoids. Scale bar represents 200 μm (n= 3 independent experiments; mean ± s.d.; **P< 0.01; two-tailed Student t-test).

**Supplementary Fig. 2.**
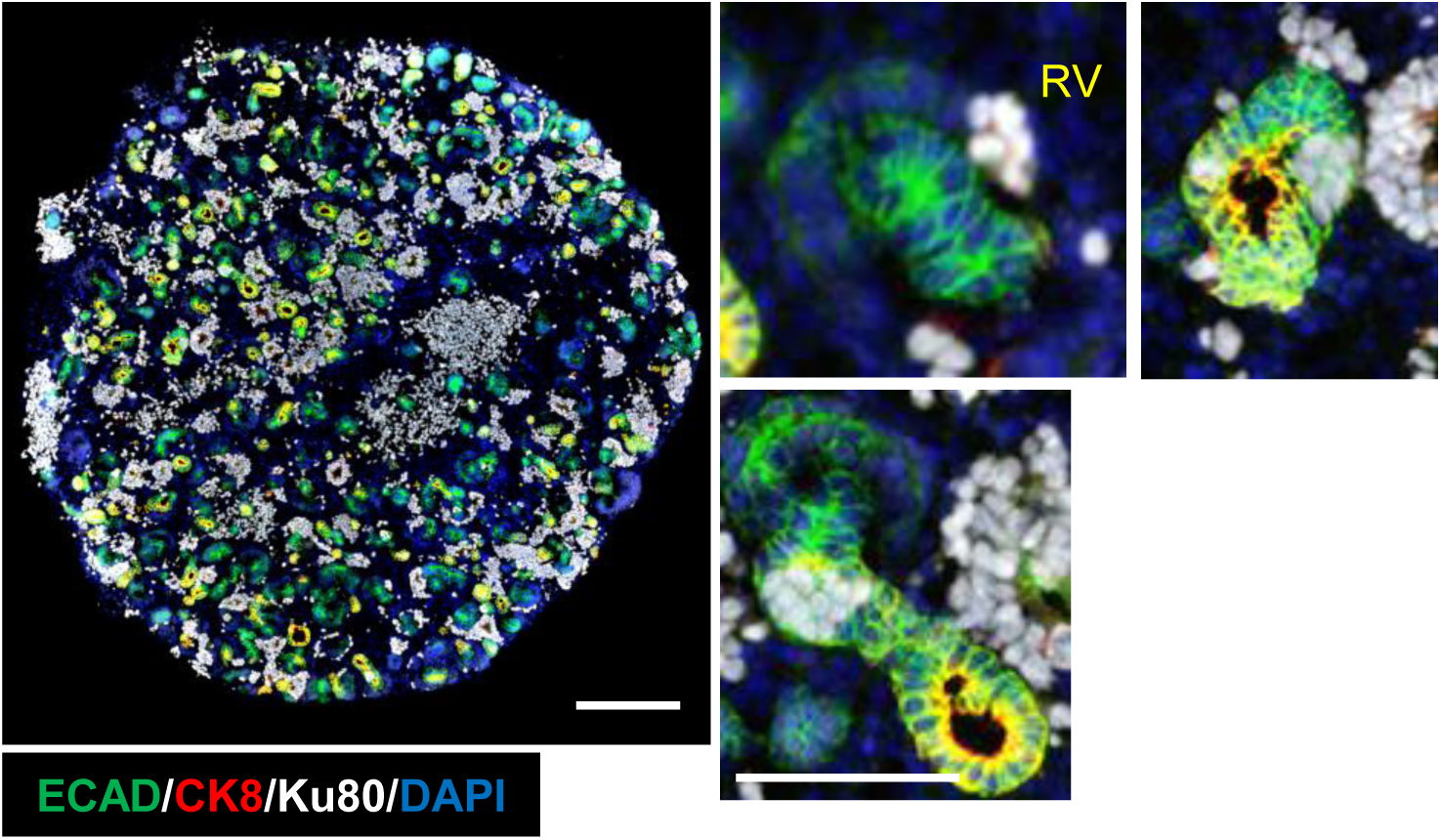
Evaluation of early-stage human-mouse chimeric renal organoids. Immunostaining images of chimeric renal organoids at Day 2 cultured in the combination of NPC_Re-agg and NPC_Mat media. Scale bar represents 200 μm.

